# A tangled tale of convergence and divergence: archaeal chromosomal proteins and Chromo-like domains in bacteria and eukaryotes

**DOI:** 10.1101/217323

**Authors:** Gurmeet Kaur, Lakshminarayan M. Iyer, Srikrishna Subramanian, L. Aravind

## Abstract

The Chromo-like superfamily of SH3-fold β-barrel domains recognize epigenetic marks in eukaryotic proteins. Their provenance has been placed either in archaea, based on apparent structural similarity to chromatin-compacting Sul7d and Cren7 proteins, or in bacteria based on the presence of sequence homologs. Using sequence and structural evidence we establish that the archaeal Cren7/Sul7 proteins emerged from a zinc ribbon (ZnR) ancestor. Further, we show that the ancestral eukaryotic Chromo-like domains evolved from bacterial precursors acquired from early endosymbioses, which already possessed an aromatic cage for recognition of modified amino-groups. These bacterial versions are part of a radiation of secreted SH3-fold domains, which spawned both chromo-like domains and classical SH3 domains in the context of peptide-recognition in the peptidoglycan. This establishes that Cren7/Sul7 converged to a "SH3”-like state from a ZnR precursor via the loss of metal-chelation and acquisition of stronger hydrophobic interactions; it is unlikely to have participated in the evolution of the chromo-like domains. We show that archaea possess several Cren7/Sul7-related proteins with intact Zn-chelating ligands, which we predict to play previously unstudied roles in cell-division comparable to the PRC barrel.

---

Three-dimensional structures or folds of proteins are more evolutionarily robust than their sequences (Murzin, 1998; Murzin et al., 1995; Schwede and Peitsch, 2008). In the absence of statistically-significant sequence similarity, structural equivalence can be used to assess evolutionary relatedness (Murzin, 1998; Orengo et al., 2001; Swindells et al., 1998). However, the evidence can, in some instances, be equivocal regarding structural convergence versus divergence: the moot point in these cases is whether the structural similarity in folds in question is a signal of a divergent origin from a common ancestor or independent convergence to a common scaffold (Doolittle, 1994; Krishna and Grishin, 2004; Lupas et al., 2001; Murzin, 1998; Zhang et al., 2014). Automated sequence- and structure-similarity search tools, though widely used for gauging relatedness among proteins, are often of limited utility in these cases. Tracing the correct relationships demands careful case-by-case analysis (Grishin, 2000; Orengo et al., 2001) and on multiple occasions, has helped untangle convergence from extreme divergence, which had otherwise eluded automated similarity search tools (Alva et al., 2008; Anantharaman and Aravind, 2004; Andreeva, 2012; Grishin, 2001a, b; Murzin, 2008; Roessler et al., 2008). In this work, we present such a case regarding the SH3 fold and certain zinc ribbons (ZnRs), with bearing on the function and evolution of key domains involved in chromatin structure and chromosome segregation in archaea, recognition of epigenetic marks in eukaryotes, and bacterial cell-wall dynamics.

The <u>S</u>rc <u>h</u>omology <u>3</u> (SH3) is a small β-barrel domain, comprised of five or six β-strands that are tightly packed into two orthogonal β-sheets (Kuriyan and Cowburn, 1993). The eponymous SH3 domains are involved in eukaryotic signaling pathways where they mediate protein-protein interactions by binding proline-rich peptide sequences via a conserved cluster of aromatic residues (Cohen et al., 1995; Kuriyan and Cowburn, 1993; Pawson, 1995). The discovery of bacterial homologs of the SH3 domain presented an interesting contrast as they were primarily found as extracellular domains in periplasmic or cell-wall associate proteins (Anantharaman and Aravind, 2003; Ponting et al., 1999; Xu et al., 2015b). Members of the larger SH3-like β-barrel fold include a vast collection of superfamilies found in diverse biological functional contexts. The best characterized of them are implicated in a variety of key protein-protein interactions via recognition of short peptide motifs (Kishan and Agrawal, 2005). SH3-like β-barrels also mediate interactions with nucleic acids (Dalgarno et al., 1997; Kishan and Agrawal, 2005). For example, the PAZ (<u>P</u>iwi <u>A</u>rgonaut and <u>Z</u>wille) SH3-like β-barrel domain, found in the Piwi and the Dicer proteins in the RNAi system interact with RNA (Burroughs et al., 2014; Lingel et al., 2003; Yan et al., 2003). Likewise, other families with the SH3 fold such as the CarD (Subramanian et al., 2000), Chromo (Bouazoune et al., 2002), TUDOR domains (Charier et al., 2004; Gong et al., 2014) have been shown to bind DNA in a number of proteins.

In recent years, it has become clear that in eukaryotes a large superfamily of domains with the SH3 fold plays a key role in recognition of short peptide-motifs, especially those with covalently modified side-chains in chromatin (chiefly histones) and RNA-processing proteins. This is the Chromo-like superfamily, which based on sequence and structure comparisons was shown to contain four major clades, including the classical Chromo (<u>Chro</u>matin organization <u>m</u>odifier) domains, BAM/BAH, BMB/PWWP and Tudor-like domains (Iyer et al., 2008; Jones et al., 2000; Koonin et al., 1995). The Tudor-like assemblage further includes the Tudor, MBT (malignant brain tumor), Agenet, DUF3590, DUF1325, RAD53BP, Tudor-knot, AuxRF(PF06507) and the MORC-C terminal domains (Aravind et al., 2011; Maurer-Stroh et al., 2003) (PFAM clan CL0049). The conserved core of the Chromo-like domains includes a SH3-like β-barrel with 5 strands that is often capped by a C-terminal helix (Jones et al., 2000; Koonin et al., 1995). They share a broadly conserved mode of interaction with peptides, specifically recognizing covalent modifications of positively charged side-chains via cation-π interactions with conserved aromatic residues (Xu et al., 2015a). While most members bind peptides with methylated lysines, the members of the Tudor-like clade specialize in binding peptides with methylated arginines.

These covalent modifications along with the Chromo-like domains that bind them are defining features of all eukaryotes, which set them apart from prokaryotes. Hence, understanding the provenance of eukaryotes depends on a proper explanation for their origins in their stem lineage. Sequence and structure comparison studies have proposed two distinct possibilities for their origins. In the first, based on structural similarity, an evolutionary relationship was proposed between the eukaryotic Chromo-like domains and the SH3-like fold of archaeal Sul7d like chromatin compaction proteins (Ball et al., 1997). These in turn are structurally and functionally related to the major pan-crenarchaeal Cren7 protein, also involved in DNA compaction and supercoiling (Guo et al., 2008; Zhang et al., 2010). Thus, in this scenario, the eukaryotic Chromo-like domains were derived from an archaeal chromatin protein in the context of chromatin function (Ball et al., 1997). In the alternative scenario, the presence of unambiguous sequence homologs of the Chromo-like domains in bacterial proteins suggest that the eukaryotic versions evolved from the bacterial precursors (Aravind et al., 2011).

To distinguish between these alternatives and better understand the function of the bacterial chromo-like domains we sought to utilize the wealth of new genomic and structural data. Using sequence and structure comparisons, we demonstrate that the Cren7/Sul7 like "SH3 fold” domains are members of the ZnR fold with certain members, including Cren7, secondarily losing their ability to chelate a zinc ion. We further shown that the Sul7 proteins most likely arose in Sulfolobales as a paralogous family of Cren7. We also show that the eukaryotic Chromo domains instead evolved from bacterial precursors as part of the radiation of SH3 fold domains in the context of bacterial cell-wall dynamics. Based on these considerations, we hypothesize that the SH3-like β-barrel architecture convergently emerged from an ancestral ZnR fold in Cren7/Sul7-like proteins.

## Results and Discussion

### Structural diversity of Zn-ribbon domains and the relationship of certain versions to the SH3-iike β-barrel fold

ZnRs are small domains that lack an extensive hydrophobic core beyond the presence of two β-hairpins, being primarily stabilized by chelation of a metal ion (Aravind and Koonin, 1999; Grishin, 2001c; Klug, 2010; Krishna et al., 2003). The metal ion is chelated by two cysteine ligands from each of the "knuckles” with the consensus motif‘CPxCG’ situated in the turn of each β-hairpin (Aravind and Koonin, 1999; Chen et al., 2000; Krishna et al., 2003). A comparative structural analysis revealed two major types of the ZnR fold, type-1 and type-2. The type-1 ZnRs have a distinct separation between the N-terminal and C-terminal β-hairpins such that no contiguous β-sheets incorporating the two knuckles or parts thereof are observed. Type-1 ZnRs can be further distinguished into two sub-types based on the presence of additional β-strand at the C-terminus. Type-1A ZnRs (Figure 1A) have a structural core made up of only two β-hairpins and are seen, for example, in the Ran binding protein family (PDB: 3CH5_B), in the pre-SET domain of few SET domain methylases (PDB: 2L9Z_A) and cytochrome c oxidase polypeptide Vb (PDB: 10CC_F). In Type-1B ZnRs, (Figure 1B), the C-terminal β-hairpin extends into an additional β-strand that forms a three-stranded β-sheet with the N-terminal β-hairpin. The type-1B ZnRs are typified by the rubredoxins (PDB: 1DX8), the ZnRs of the tRNA synthetases (PDB: 1A8H) and the ubiquitin-binding ZnRs of deubiquitinating peptidases of the UBCH family (PDB: 2GFO).

**Figure 1:**
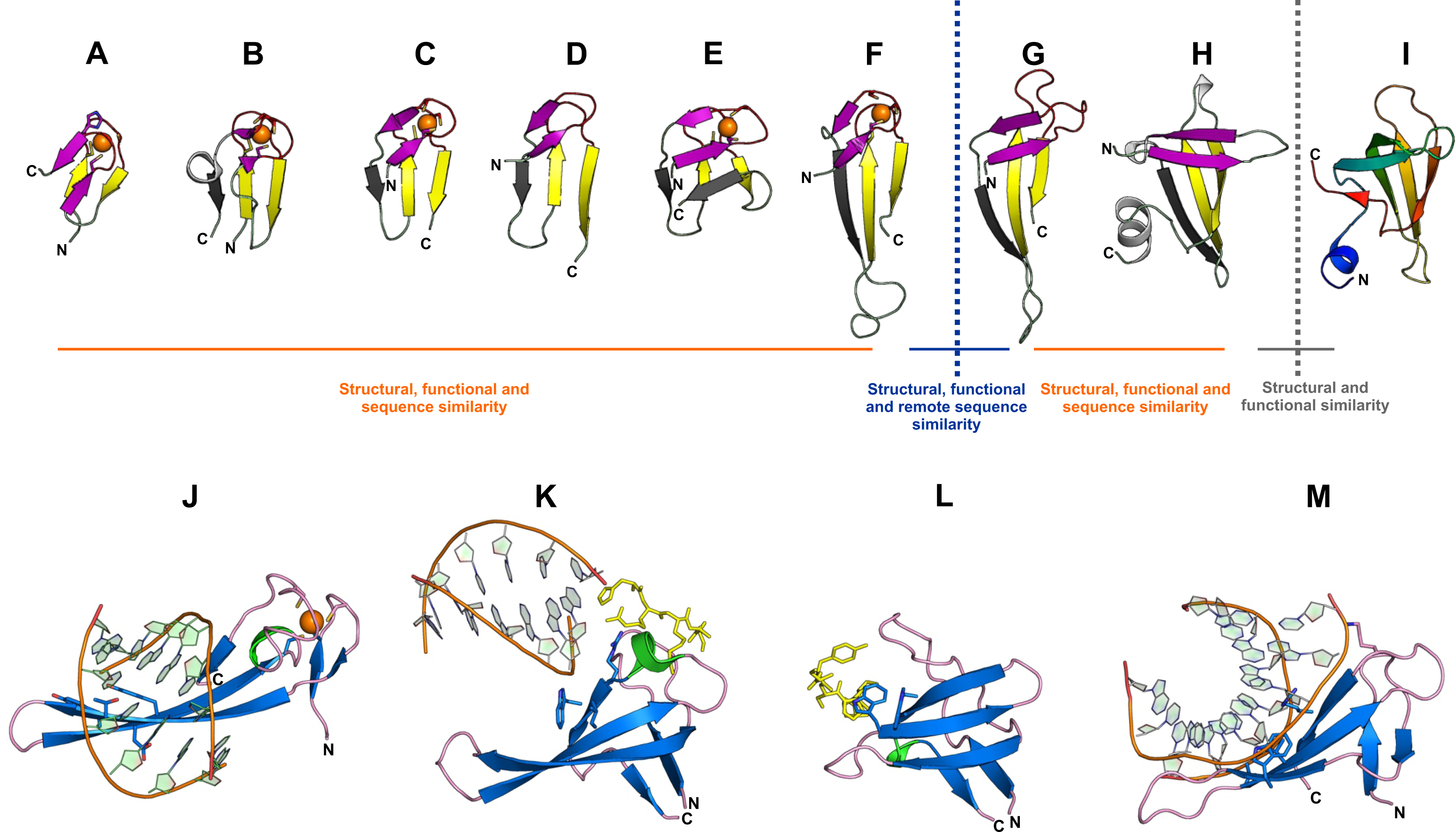
Comparative structural view of the major secondary elements (A-I) and binding interfaces (J-M) in ZnRs, SH3s and Cren7. **(A)** Type-1A ZnR (PR domain-containing protein 11, PDB: 3RAY:A) **(B)** Type-1B zinc ribbon (Rubrerythrin, PDB: 1NNQ:A) **(C)** Type-2 zinc ribbon (Archaeal exosome RNA binding protein CSL4, PDB: 2BA1:A) (D) Type-2 zinc ribbon (Uncharacterized BCR *cg1592,* PDB: 2JNY:A) (E) ZnRs with three strands on each side (Anaerobic ribonucleoside-triphosphate reductase, PDB: 1H8K:A) **(F)** Type-2 zinc finger (Transcriptional Elongation Factor SII, PDB: 1TFI:A) **(G)** Cren7 (PDB: 3KXT:A) **(H)** Sso7 (PDB: 1C8C:A) **(I)** SH3-like fold (Tudor domain-containing protein 3, PDB: 3PNW:0). **(J)** DNA bound zinc ribbon domain (putative integrase [Bacteriophage A118], PDB: 4KIS:A) (K) DNA and peptide bound chromodomain (Male specific Lethal-3, PDB: 30A6:A) (L) peptide bound SH3 domain (Tyrosine-protein kinase ABL1, PDB: 1BBZ:A) **(M)** DNA bound Cren7 (PDB: 3KXT_A). In all the panels, the Zn ion is represented by an orange sphere, side chains of zinc-chelating and other functionally-important amino acids are represented as sticks, bound peptides are yellow colored stick, the sugar-phosphate backbone of DNA is orange with nucleotides in green. Color scheme for **(A-I):** For type-2 zinc ribbons, the N-terminal (β-hairpin is colored purple, the third (3-strand is colored gray, the zinc knuckles are colored red and the C-terminal β-hairpin is colored yellow. The equivalent β-strands in type-2, −1A and −1B are colored alike. The Tudor domain is colored in a gradient of blue to red from the N-to the C-terminal. In panels **(J-M):** All β-strands are colored in blue, all loops in pink and α-helices in green.

In type-2 ZnRs the N-terminal β-hairpin is extended into a β-strand that packs with C-terminal β-hairpin via hydrogen bonds forming a three-stranded β-meander (Figure 1C-F). Type-2 ZnRs tend to be prevalent in nucleic acid-binding proteins (PDB: 1TFI, 1L10) and in a strand-swapped version in the DNA-binding Ku and MarR family ZnR domains (PDB: 1JEY, 2F2E) (Aravind and Koonin, 1999; Kaur and Subramanian, 2015; Krishna and Aravind, 2010). Despite these structural differences the two classes of ZnRs are likely to be related because they possess a common four stranded core, conserved geometry of the Zn-chelating residues and can be structurally superimposed.

Interestingly, while developing this classification of ZnRs, we noticed a consistent structural similarity between type-2 and SH3 domains (Figure 1I). For example, a DALI (Holm and Sander, 1995) search initiated with the ZnRs from the RNA polymerase subunit RBP9 (PDB: 1QYP_A), in addition to retrieving various ZnRs (e.g. Ribonuclease P protein component 4, PDB: 1X0T_A, Z-score=3.9, RMSD=1.2Å, lali=38; zinc finger protein ZPR1, PDB: 2QKD_A, Z-score=3.3Å, RMSD=3.3Å, lali=40), recovered several distinct SH3-like β-barrel domains albeit with lower Z-scores such as the classic SH3 domains (e.g. Myosin VI, PDB: 2VB6_A, Z-score=2.2, RMSD=1.8υ, lali=33; Rho guanine exchange factor 16, PDB: 1X6B, Z-score=2.2, RMSD=5.lυ, lali=43), the Tudor domain (e.g. Survival motor neuron protein, PDB: 4A4E_A, Z-score=2.5, RMSD=2.9Å, lali=41) and the Chromo domain (e.g. Mortality factor 4-like protein 1, PDB: 2F5K_F, Z-score=2.0, RMSD=2.6Å, lali=39). Several SH3-like β-barrel folds were also retrieved in structural searches initiated with the zinc ribbon domains of peptide:N-glycanase (PDB: 1X3W_A), ORF131 of *Pyrobaculum* spherical virus (PDB: 2X5C_A), anaerobic ribonucleotide-triphosphate reductase (PDB: 1HK8_A) and lysyl-tRNA synthetase (PDB: 1IRX_A). Further, visual inspection confirmed this relationship, indicating a topological similarity in arrangement of the β-strands in type-2 ZnRs and certain members of the SH3-like β-barrel fold.

### Type-2 ZnRs and SH3-fold domains show comparable ligand-binding interfaces

The detection of this relationship between type-2 ZnRs and SH3-fold domains led us to further compare the ligand-binding interfaces of members of the two folds. In type-2 ZnRs the ligand-binding surface is formed by the three-stranded β-sheet typical of these domains (Krishna et al., 2003). The interacting residues often emanate from strands β3 and β4 and frequently have aromatic or charged side chains (Figure 1J). For example, this interface is used to contact nucleic acids by ZnRs in proteins TFIIS, TBP N-terminal domain, Ku, RBP9 and the RNaseP Rpp21 (Awrey et al., 1998; Olmsted et al., 1998); (Amero et al., 2008). Interestingly, while Chromo-like domains bind methylated peptides mainly via the open region of the barrel, their nucleic-acid-binding interface is structurally comparable to the ligand-binding interface of the Type-2 ZnRs. (Figure 1K). The classical SH3 domains use the same interface as the Chromo-like domains to mediate protein-protein interactions through a tryptophan residue which interacts with the proline-rich peptide substrate (Cohen et al., 1995; Pawson, 1995) (Figure 1L). The position of this tryptophan corresponds to that of the nucleic acid-interacting residues in ZnRs. These observations suggest that in addition to the structural similarities, at least one binding interface of the SH3-fold domains is similar that of the type-2 ZnRs. This analysis also led us to the archaeal chromosomal proteins Cren7 (PDB: 3KXT; Figure 1G) and Sac7d/Sul7 (PDB: 1WD0; Figure 1H) proteins, which have been classified with the Chromo-like SH3 fold domains, and have even been proposed as their precursors. In these proteins too, the residues responsible for DNA-binding are mainly contributed by the region of the triple-stranded β-sheet (β3-β4-β5) (Figure 1M) (Guo et al., 2008; Robinson et al., 1998);(Guo et al., 2008; Zhang et al., 2015) which presents a clear parallel to the type-2 ZnRs.

### The origin of Cren7 and Sul7 proteins from ZnRs and their diversiffcation in archaea

The above observations hinted that that Cren7 with structural features of both SH3-fold and ZnR domains might help better understand the similarities between the two folds. Consistent with the findings of Guo *et al,* 2008, DALI searches initiated with Cren7 protein (PDB: 3KXT_A), recovered multiple SH3-like fold domains such as Chromo domains, Tudor domains and myosin S1 fragment SH3 domains (Guo et al., 2008). These searches also recovered the Sul7 protein (Guo et al., 2008; Zhang et al., 2010) (PDB: 1WVL_A), which is believed to be structurally and functionally related to Cren7 as one of the hits (PDB: 1WVL_A, Z-score: 3.3, RMSD=3.1 Å lali=46). Concurrently, the search also retrieved hits to ZnRs, such as those in sarcosine oxidase delta subunit (PDB: 1VRQ_D, Z-score=3.0, RMSD=2.8Å, lali=44), PNGase (PDB: 3ESW_A, Z-score=2.8, RMSD=2.6Å, lali=42), zinc finger protein ZPR1 (PDB: 2QKD_A, Z-score=2.4, RMSD=3.4Å, lali=44), peptide:N-glycanase (PDB: 1X3W_A, Z-score=2.2, RMSD=3.1Å, lali=43) and 50S ribosomal protein L44E (PDB: 1Q81_4, Z-score=2.0, RMSD=3.2Å, lali=45). Manual structural superimposition of Cren7 (PDB: 3KXT_A) and ZnRs (eg. sarcosine oxidase delta subunit, PDB: 1VRQ_D) perfectly aligned all secondary structure elements of the Cren7 β-barrel onto the core of the ZnR fold (RMSD= 1.5 Å over 35 pairs of backbone C_α_ atoms), where the turns of β1/β2 and of β4/β5 of Cren7 recapitulate the position of knuckles in ZnRs. Similar results were obtained by automated pairwise structural alignment of Cren7 (PDB: 3KXT_A) and ZnRs (eg. sarcosine oxidase delta subunit, PDB: 1VRQ_D) using TM-align (Zhang and Skolnick, 2005) and Fr-TM-align (Pandit and Skolnick, 2008) which gave a TM score of 0.52 (normalised by the length of 3KXT_A), indicating a 'fold level' similarity between the two.

To investigate if this structural relationship to ZnRs also extends to sequence similarity, we initiated iterative sequence similarity searches with *S. solfataricus* Cren7 (PDB: 3KXT_A) using the PSI-BLAST (Altschul et al., 1997) and the JACKHMMER (Finn et al., 2011) programs. Interestingly, these searches recovered multiple orthologous sequences crenarchaeota and bathyarchaeota that contained two to four cysteine residues at positions corresponding to turns between strands β1/β2 and β4/β5 (eg. *Ignicoccus hospitalis,* WP_011998075.1) (Figure 2A). The positions of these cysteines suggest that when four of them are present they are likely to chelate a Zn ion. Moreover, we found another group of Cren7 homologs from several crenarchaea which contained a Cren7 domain as part of a larger multidomain protein (see below). These Cren7 domains were characterized by the presence of all four expected Zn-chelating cysteines. Additionally, our searches also retrieved a distinct paralogous family of crenarchaeal proteins (Cren7Znr family; e.g. APE_2454.1 from *Aeropyrum pernix,* BAA81469.2) typified by an extended C-terminal region, most of whose members possess intact Zn-chelating cysteines (Figure 2A). Profile-profile sequence similarity searches (Soding et al., 2005) initiated with the Cren7 and Cren7ZnR domains consistently retrieved ZnRs. For example, HHPRED searches with *I. hospitalis (WP _011998075.1)* found matches to many ZnR proteins such as probable lysine biosynthesis protein LysX/ArgZ (PDB: 5K2M_E, E-value= 0.0005), ZitP pilus assembly/motility regulator (PDB: 2NB9_A, E-value= 0.0054) and transcriptional regulator MqsA (PDB: 309X_A, E-value= 0.091). Further, even among the Cren7Znr proteins some representatives show loss of one of the Zn-chelating cysteines exactly paralleling the situation observed among the classic Cren7 proteins (Figure 2A).

**Figure 2.**
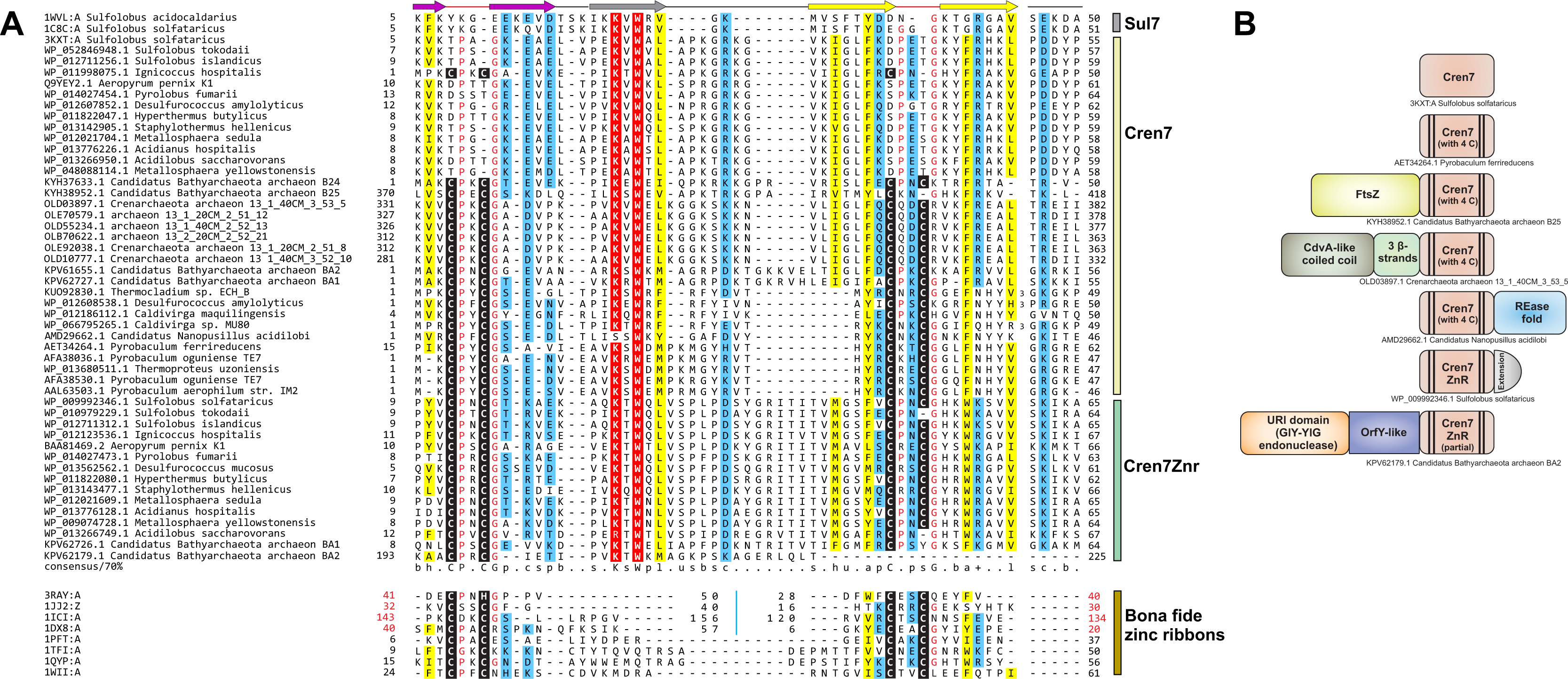
Multiple sequence alignment (MSA) and domain architectures of Cren7 proteins. (A) Structure-based MSA of representative sequences of Sul7, Cren7, Cren7Znr and bona fide ZnRs. Accession number or the PDB id, and the alignment range are indicated for each sequence. For Sul7, Cren7 and Cren7Znr proteins, the organism names are also mentioned after the accession number/PDB. Zn-chelating residues are boxed in black and small amino acids (Gly, Pro) just after the Zn-binding ligands are colored in red. At positions with a 70% consensus, charged or polar residues are highlighted in blue, and hydrophobic, apolar and aliphatic residues are in yellow. Some insertions are not displayed for clarity, and the number of omitted residues is indicated. Regions of circular permutation in ZnRs are separated by a blue ‘| ’ mark and the sequence numbers of the permuted ZnRs are shown in red. The consensus secondary structures are depicted above the alignment. **(B) Domain architectures of Cren7 and Cren7Znr proteins.** Representative sequence identifier with organism name are mention below individual architectures.

Thus, multiple lines of evidence support that Cren7-like proteins (PDB: 3KXT_A) are ZnRs, with secondary loss of Zn-chelating residues in some versions such as the *S. solfataricus* Cren7. Although the Sul7 proteins were not retrieved in these sequence searches, they show several features that suggests their derivation from Cren7 proteins. 1) They are so far only found in a limited number of crenarchaeal lineages (*Sulfolobus, Acidanus* and *Metallosphaera)* unlike Cren7, which is conserved across crenarchaeota and the bathyarchaeota (Evans et al., 2015). 2) They share a common DNA-compaction function in crenarchaeal chromatin (Guo et al., 2008; Zhang et al., 2015; Zhang et al., 2010), which is mediated via a similar protein-DNA interface, with positionally- and chemically-equivalent residues involved in DNA contact. 3) Experiments to make the Cren7 protein more "Sul7d-like” by mutating the loop between strands β3/β4 results in an exact superposition of the DNA-binding interface (Guo et al., 2008; Zhang et al., 2015; Zhang et al., 2010).

Presence of Cren7/Sul7 proteins only in crenarchaeota and bathyarchaeota, together with their provenance from ZnRs, which we establish above supports the following evolutionary scenario: the ancestral Cren7 proteins likely arose from a DNA-binding ZnR of which several are found in the pan-archaeo-eukaryotic transcription apparatus (Aravind and Koonin, 1999). Consistent with this, we have found versions of Cren7, which still retain the ancestral Zn-chelating residues. The strengthening of the hydrophobic core due to interactions in the strand regions appear to have facilitated the loss of Zn-chelation in more than one version of the family. This appears to have concomitantly supported the emergence of a more barrel-like geometry that converged to a SH3-like state. This was followed by the emergence of Sul7 only in Sulfolobaceae as a specialized DNA-packaging protein from a Cren7-like protein that had already lost its Zn-chelating residues.

Our recovery of novel members of the Cren7-like family suggests that they underwent functional diversification in the archaeal lineages that possess them. Notably, in one group of these proteins, Cren7 is the C-terminal domain of much larger protein (e.g. OLD03897.1) where it is combined with an N-terminal CdvA-like coiled coil domain and a central small 3 β-stranded domain. In crenarchaea the CdvA-like coiled-coil domain proteins are components of the cell-division system and form filamentous double-helical complexes with DNA (Moriscot et al., 2011). In bathyarchaea, we found a fusion of the Cren7 domain to an N-terminal FtsZ domain (KYH38952.1), a key component of the cell-division apparatus related to the tubulin-like cytoskeletal proteins. These architectures suggest that representatives of the Cren7 family play a specific role in cell-division, probably in anchoring the DNA. All Cren7Znr proteins contain a conserved C-terminal motif with an absolutely conserved acidic residue beyond the core Cren7 domain, which is likely to adopt an extended conformation. It is possible that this region also helps in specific interactions with cell-division components. We also found an instance of bathyarchaeal Cren7 (KPV62179.1) fused to the URI domain (GIY-YIG endonuclease) and an uncharacterized enzymatic domain of the OrfY-like superfamily (Kryshtafovych et al., 2014), and a nanoarchaeal Cren7 (AMD29662.1) with a C-terminal REase fold nuclease domain (Figure 2B).

### Bacterial diversification of Chromo-like SH3 fold domains

The above rooting of the provenance of Cren7/Sul7-like proteins within the archaeal radiation of ZnRs and evidence for convergent acquisition of the SH3-like barrel morphology questions the evolutionary relationship of these archaeal chromosomal proteins and the Chromo-like superfamily of SH3 fold, especially given the previous identification of bacterial versions of the latter (Aravind et al., 2011). To better understand the origins of the bacterial Chromo-like domains, we ran iterative profile searches from the versions we had previously detected in bacteria (Aravind et al., 2011). Our current searches, greatly extended the phyletic spread of bacterial chromo-like domains, retrieving them from several different lineages such as proteobacteria (mainly α and δ), cyanobacteria, Thermus/Deinococcus, bacteroidetes, planctomycetes, and spirochaetes and in rare cases in euryarchaea. Firmicutes and actinobacteria showed a strong under-representation of this domain (see **supplementary material**). A multiple sequence alignment of the bacterial Chromo-like domains with various eukaryotic versions revealed that the bacterial homologs strongly conserve at least a subset of the aromatic residues corresponding to the aromatic cage involved in ligand-binding (Figure 3A, B) (Aravind et al., 2011; Kim et al., 2010; Nielsen et al., 2002). Given that these residues are central to the recognition of methylated Ɛ-amino groups of lysine in the bound peptide in eukaryotic Chromo-like domains (Figure 3C), we posit a similar binding capacity for the bacterial versions. Across eukaryotic Chromo-like domains, the Tudor assemblage strongly conserves the residues forming the aromatic cage, whereas the classical Chromo-domains shows some variability in these residues (Figure 4). This suggests that the bacterial versions are closer to the Tudor-like Chromo domains and that the Tudor-like versions are likely to be closer to the ancestral mode of binding peptides. The plausible phylogenetic relationship shared by the various Chromo-like domains is depicted in figure 3D.

**Figure 3.**
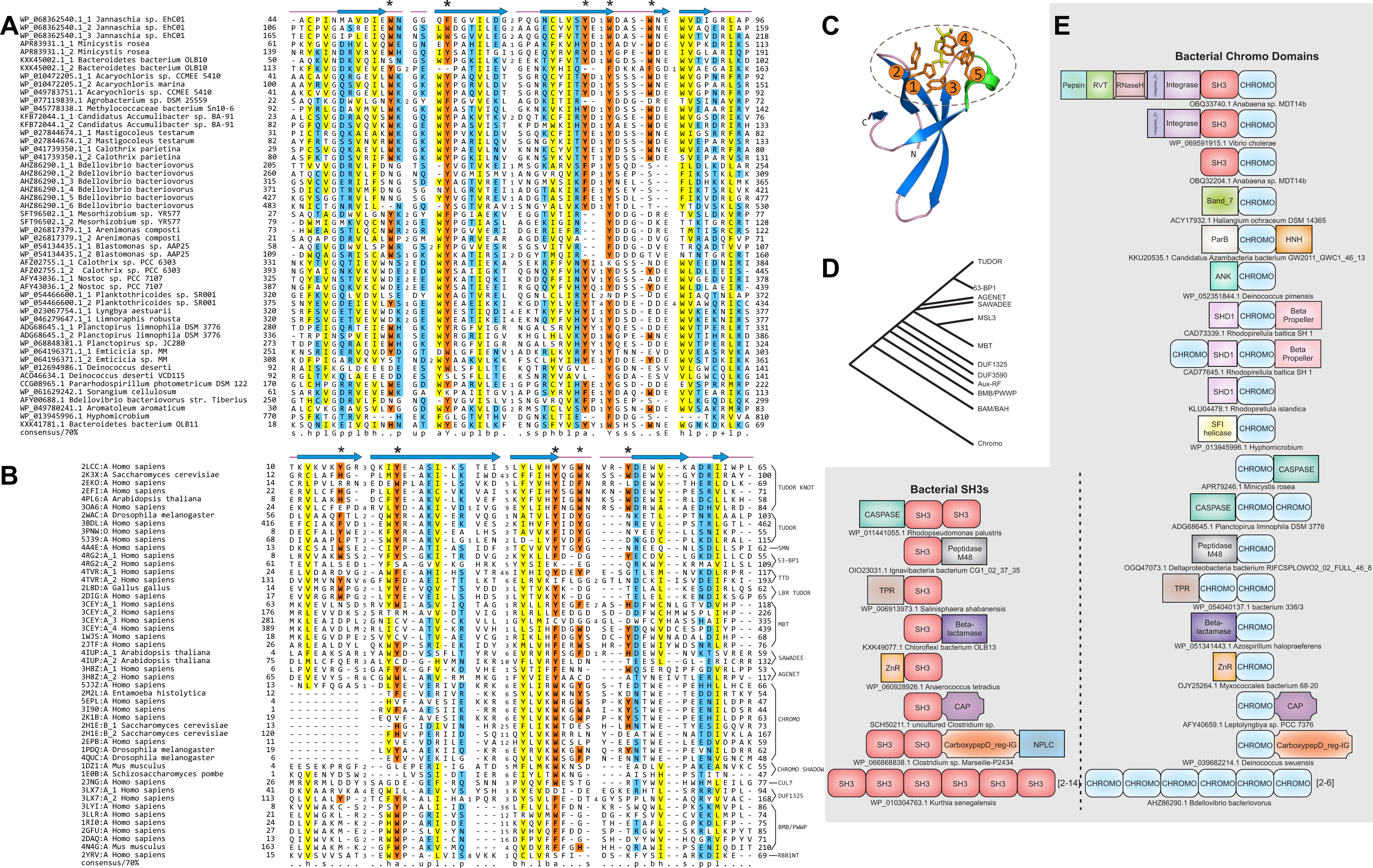
Multiple sequence alignment (MSA), probable phylogeny and domain architectures of Chromo-like domains. (A) MSA of representative bacterial Chromo-like domains. (B) Structure-based MSA of representative eukaryotic chromo-like domains. **Coloring scheme for panel (A) and (B)** follows figure **2(A)**. The amino acids at positions that are likely to be involved in forming the peptide-binding aromatic cage are highlighted in orange and marked with the asterisk above the MSA. **(C) View of the peptide-binding aromatic cage of** *Arabidopsis* Morf **Related Gene (MRG) group protein MRG2 Chromo domain (PDB: 4PL6**). The aromatic cage is marked by a dashed eclipse and the aromatic residue positions from the N- to C-terminus are marked by numbers 1-5. **(D) Probable phylogenetic relationships between the various Chromo-like domains. (E) Domain architectures of bacterial Chromo domains.** Parallel architectures among bacterial Chromo and bacterial SH3 domains are grouped at the bottom.

**Figure 4.**
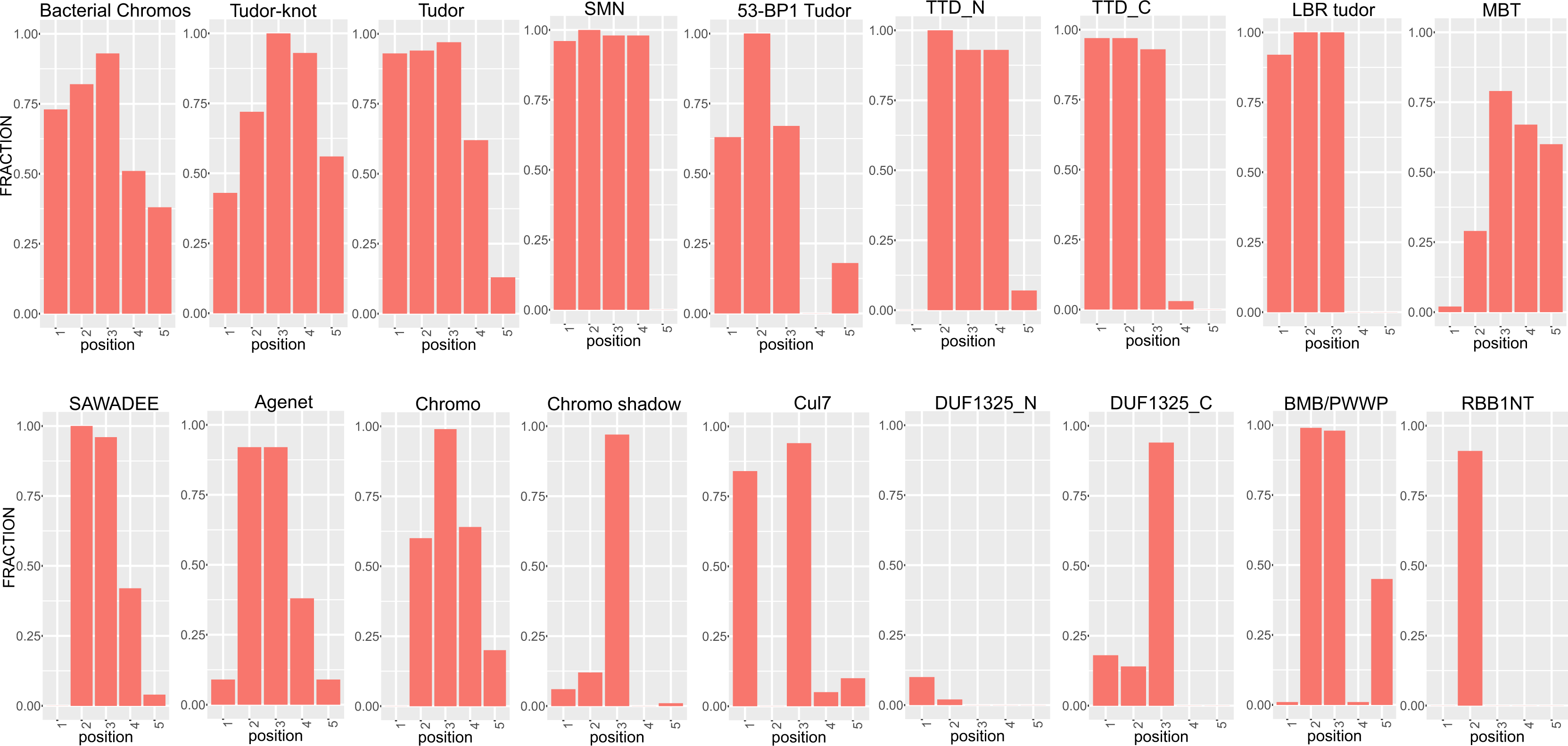
Comparative graphical view of the conservation of aromatic residues at the five positions that make up the peptide-binding aromatic cage in Chromo-like domains. Individual graphs are labelled on the top by the family they represent. The X-axis represents positions corresponding to those shown in figure 3(C). The Y-axis represents the fraction of aromatic amino acids (Y, W, F, H) at those positions in the respective families.

We observed that the bacterial Chromo-like domains, in contrast to their eukaryotic counterparts, are consistently present in secreted proteins across the diverse lineages containing them. A systematic analysis of their domain architectures again revealed contexts unlike any seen in eukaryotes (Figure 3E): in addition to being found in tandem repeats (2-6 domains per protein), one of the most common linkages of the secreted bacterial Chromo-like domains is with the caspase-like peptidase domain. This architecture is found in plantomycetes, cyanobacteria, bacteriodetes and chloroflexi. Notably, less-frequent but parallel fusions are also observed with other extracellular enzymatic domains namely a zincin-like metallopeptidase and a metallo-β-lactamase domain which is predicted to function as a nuclease (Figure 3E). Additionally, these Chromo-like domains are also combined in extracellular proteins with several other non-catalytic domains, such as another SH3-fold domain the Slap homology domain 1 (SHD1), WD40-like β-propeller domains, the EF-hand, and the Ig-fold carboxypeptidase regulatory domain TPR repeats. Interestingly, parallel domain architectures with multiple tandem domains and fusions to the caspase, zincin-like metallopeptidase, WD40 β-propellers, metallo-β-lactamase, the carboxypeptidase regulatory and TPR repeat domains are seen for bacterial representatives of classical SH3 domains (Figure 3B) (Anantharaman and Aravind, 2003; Ponting et al., 1999), again in contrast to their strictly intracellular eukaryotic counterparts (Pawson, 1995). These bacterial SH3 domain proteins are distributed across a much wider phyletic range of taxa compared to the Chromo-like domains. Recent studies have shown that secreted bacterial SH3 domains are likely involved in binding peptides in the peptidoglycan cell wall ((Anantharaman and Aravind, 2003; Ponting et al., 1999; Xu et al., 2015b). These parallels suggest that the bacterial Chromo-like domains might function similarly to the bacterial SH3 in binding-specific peptides. However, they are likely to possess specificity for those containing methylated lysine-like moieties, in the murein or extracellular matrix peptides in bacteria with Gram-negative cell walls (given their near absence in firmicutes and actinobacteria).

In addition to the above architectures, we also observed two unusual lateral transfers of the eukaryotic Chromo domains to bacteria: 1) a Ty-3 family retrotransposon, which is commonly found in fungi is also found in multiple copies in *Anabaena* (for example, Accession no. OBQ33740.1, from *An*abaena sp. MDT14b). Here the Chromo domain is fused C-terminal to the polyprotein containing pepsin, reverse transcriptase, RNase H, integrase and SH3 domains. Given the association of certain fungi with cyanobacteria (e.g. example, in cyanolichens) this probably represents a transfer facilitated by such an association. 2) The other case is found in a single species, where a eukaryotic chromodomain is inserted into a mobile element of a bacterium with a ParB and HNH domains (accession: KKU20535.1, Azambacteria bacterium GW2011_GWC1_46_13) (see **Supplementary material**).

## Conclusions

Small metal-chelating domains, such as ZnRs, likely originated with a relatively simple stabilizing core in the form of the Zn-chelating center. The ancestral ZnRs themselves could have emerged from a pair of small metal-stabilized, bi-cysteine, knuckle-like motifs that existed independently (for example, similar to the minimal versions seen in Rad50 zinc-hook, PDB: 1L8D; RNase E zinc-link domain, PDB: 2VMK). More structured ZnRs with β-hairpins developing around these Zn-knuckles at their turns (Figure 1A-F) likely evolved from such versions and acquired the ability to exist as independent domains. These simplest versions of ZnRs might have resembled the type-1 A scaffold as they have separate β-hairpins with the two metal chelating residue pairs (Figure 1A). This provided a platform for considerable evolutionary variability and structural innovation with augmentation and/or supplanting of the original Zn-center by emergence of further stabilizing hydrogen-bonding networks and hydrophobic cores in the form of new, more ordered secondary structure elements (Aravind et al., 2006; Arnold and Zhang, 1994; Salgado et al., 2010; Zhang et al., 2014). Such developments are seen in the form of the type-1 B ZnRs, which developed an additional β-strand and finally the type-2 ZnRs, which appear to be related to the type-1B versions via a circular permutation. Here, the additional β-strand appears to have been further incorporated into the core of the fold to form a contiguous β-sheet (Figure 1C-F). Indeed, loss of metal-ion chelation in such domains can accompany the development of alternative stabilizing hydrophobic cores, which might be the progenitor of a distinct protein fold (Aravind and Koonin, 2000; Krishna and Aravind, 2010). This could be subject to further modifications via duplications and/or circular permutations (Burroughs et al., 2011; Krishna et al., 2003)

In this study, we capture the evolutionary stages in one such transformation: the Cren7 domain previously considered a SH3 fold β-barrel actually emerged from a metal-binding ZnR domain. We find evidence that such convergence between ZnRs and SH3 fold domains might have happened independently more than once. For example, the segment-swapped Ku-bridge domain and the similar C-terminal all-β domain of the MarR-like transcription factors (labelled as SH3-like in SCOP and SC0P2; SCOP identifier 140307) resemble segment-swapped SH3-like folds; however, sequence analysis clearly indicated the provenance of these domains from type-2 ZnRs (Kaur and Subramanian, 2015; Krishna and Aravind, 2010). We also observed that the type*-2* ZnR at C-terminus of the nicotinate phosphoribosyltransferase converged to a SH3-like fold with the Zn-chelating sites lost alongside the evolution of compensatory hydrophobic interactions (**Supplementary Figure 1**). Further, this convergence can also extend to the substrate-binding mode of certain ZnRs and the SH3 fold domains (Figure 1J-M). This raises the possibility that similar transitions might have spawned SH3-like fold domains on other occasions but cannot be currently confirmed using sequence-based methods. Notably, both ZnRs and SH3 fold domains are found in proteins highly conserved across life, such as the ribosomal subunits (Anantharaman et al., 2002). Hence, it is possible that the ancient representatives of the two folds also share an ultimately divergent relationship in a very early period of the evolution of protein universe prior to the last universal common ancestor (LUCA), with the SH3 fold emerging via loss of Zn-chelation from a ZnR via acquisition of stronger hydrophobic interactions.

Our findings help clarify the origins and new functions of key domains in chromosomal proteins. Compaction of genomic DNA into the limited intracellular cell is a universal solution which has selected for multiple solutions across cellular life. In asgardarchaea (believed by many to be the closest sister group of eukaryotes), eukaryotes and euryarchaea this function is carried out by the α-helical histone fold proteins and in bacteria by the Hu/IHF superfamily of proteins (Burroughs et al., 2017; Dillon and Dorman, 2010; Luger et al., 1997; Reeve et al., 2004). However, in crenarchaeaota and least certain bathyarchaeota, the ancestral histones appear to have been displaced by a heterogeneity of chromatin-compacting proteins, namely Cren7, Sul7d and CC1 (Zhang et al., 2012). Based on structural considerations it was earlier suggested that these archaeal chromosomal proteins might have an evolutionary a relationship to the SH3-fold domains of the chromo-like superfamily, which are a hallmark of eukaryotic chromatin proteins (Ball et al., 1997). Here, we present a clear evolutionary scenario for the convergent origin of a SH3-like morphology for Cren7/Sul7d-like proteins from ZnRs. We also present evidence for a functional radiation of these ZnRs in crenarchaeaota and bathyarchaeota with potential roles in cell-division comparable to the CdvA-like proteins with the PRC-barrel domain (Anantharaman and Aravind, 2002; Moriscot et al., 2011).

On the other hand, we present evidence that eukaryotic Chromo-like domains have no close relationship to Cren7-like archaeal chromosomal proteins. While there are several SH3 fold β-barrels, eukaryotic Chromo-like domains specifically share a peptide-binding function with the classical SH3 domains. We show that both these superfamilies are present in bacterial extracellular proteins and based on the evidence from the bacterial SH3 domains, we posit that the bacterial Chromo-like domains too bind peptides in the peptidoglycan or in the periplasm. Further, sequence analysis suggests the bacterial Chromo-like domains likely acquired specificity for methylated side-chains of basic amino acids even in these bacterial versions. These observations suggest that the two SH3 fold superfamilies initially diverged and radiated primarily in the context of binding different peptides in the bacterial peptidoglycan context. Notably, both these show parallel domain architectures (Figure 3E) and are often coupled with peptidase or nuclease domains which might play role in the degradation of proteins and extracellular DNA found in bacterial extracellular-matrices. Thus, we predict that both domains might help anchor enzymes regulating the dynamics of cell-cell interactions in bacteria.

These observations have important implications for the origin of eukaryotes. Bacterial Chromo-like domains are found only in certain bacterial lineages, including α-proteobacteria, unlike the bacterial SH3 domain which is more widely distributed. However, Chromo-like domains are rare or absent in archaea (including asgardarchaeota). This suggests that the eukaryotes possibly acquired their Chromo-like and classical SH3 domains from the α-proteobacterial mitochondrial progenitor. Thus, their presence in the extracellular matrix might have facilitated interactions with “cytoplasmic” proteins and nucleic acids of archaeal component of the ancestral eukaryote upon endosymbiosis. Hence, we posit that this was the likely scenario that favoured their recruitment as intracellular peptide-binding domains in the ancestral eukaryote. In support of this, we can trace at least 3 paralogous families of Chromo-like domains in the LUCA(Aravind et al., 2011) - the Chromo-like superfamily was acquired by the stem eukaryote during the primary symbiogenetic event. In eukaryotes, the Chromo-like and classical SH3 domains radiated extensively due to the "opening” of entirely new niches in the form of peptide-substrates from histone tails, positively charged RNA-binding complexes and cytoskeletal proteins respectively. This appears to have gone hand-in-hand with the radiation of a diverse suite of enzymes covalently-modifying histones and other proteins (Aravind et al., 2011). Since bacterial Chromo-domains already likely bound peptides through the aromatic cage in the open mouth of the β-barrel, they were pre-adapted to binding methylated peptides in the eukaryotic chromatin niche.

We believe the Cren7-like and Chromo-like domains identified in this study might help further experimental characterization of the biochemical and biological diversity of these domains.

## Methods

The PSI-BLAST and JACKHMMER programs were used for iterative sequence profile searches against the National Center for Biotechnology Information (NCBI) non-redundant (NR) and locally clustered databases (eg. all.90: sequences clustered at 90% sequence identity) (Altschul et al., 1997; Finn et al., 2015). Additional sequence similarity searches were performed using the HHpred program (Soding et al., 2005) (against: PDB70_12Febl7, SC0P95_vl.75B and PfamA_31.0, using MSA generation method HHblits run for 5 iterations, E-value threshold of 0.001). Multiple sequence alignments were constructed using Kalign (Lassmann et al., 2009) followed by manual tweaking based on structural alignments. Sequence similarity-based clustering was performed using the BLASTCLUST program (ftp://ftp.ncbi.nih.gov/blast/documents/blastclust.htmn. by adjusting the length (L) and score (S) parameters based on need. Automated structure similarity searches were performed using the DALI server (Holm and Sander, 1995). Structures were compared and superimposed in the molecular visualization program PyMOL by manually defining equivalent regions using the pair fitting wizard. Automated pairwise structure superimposition was performed using TM-align and Fr-TM-align tools (Pandit and Skolnick, 2008; Zhang and Skolnick, 2005).

Domain architectures and other contextual information about the protein sequences were retrieved using in-house collection of PERL scripts for automation of analysis. Graphs in figure 4 were generated using multiple sequence alignments (MSAs) of representative sequences of each family in a local database with NR sequences clustered down to 90% identity by analyzing amino acid conservation at the five positions involved in the formation of aromatic age. Graphs were built using ggplot2 package in R (Wickham, 2009).

## Acknowledgements

This work was supported by the funds of the Intramural Research Program of the National Library of Medicine, USA (LA, LMI and GK) and the department of Biotechnology, India (BTISNET; GAP001, SS).

## Supplementary Data

Supplemental data is also available at: <u>ftp://ftp.ncbi.nih.gov/pub/aravind/temp/CREN/CREN.html</u>

## References

Altschul, S.F., Madden, T.L., Schaffer, A.A., Zhang, J., Zhang, Z., Miller, W., and Lipman, D.J. (1997). Gapped BLAST and PSI-BLAST: a new generation of protein database search programs. Nucleic acids research 25, 3389–3402.

Alva, V., Koretke, K.K., Coles, M., and Lupas, A.N. (2008). Cradle-loop barrels and the concept of metafolds in protein classification by natural descent. Current opinion in structural biology 18, 358–365.

Amero, C.D., Boomershine, W.P., Xu, Y., and Foster, M. (2008). Solution structure of Pyrococcus furiosus RPP21, a component of the archaeal RNase P holoenzyme, and interactions with its RPP29 protein partner. Biochemistry 47, 11704–11710.

Anantharaman, V., and Aravind, L. (2002). The PRC-barrel: a widespread, conserved domain shared by photosynthetic reaction center subunits and proteins of RNA metabolism. Genome biology 3, Research0061.

Anantharaman, V., and Aravind, L. (2003). Evolutionary history, structural features and biochemical diversity of the NlpC/P60 superfamily of enzymes. Genome biology 4, R11.

Anantharaman, V., and Aravind, L. (2004). The SHS2 module is a common structural theme in functionally diverse protein groups, like Rpb7p, FtsA, GyrI, and MTH1598/TM1083 superfamilies. Proteins 56, 795–807.

Anantharaman, V., Koonin, E.V., and Aravind, L. (2002). Comparative genomics and evolution of proteins involved in RNA metabolism. Nucleic acids research 30, 1427–1464.

Andreeva, A. (2012). Classification of proteins: available structural space for molecular modeling. Methods Mol Biol 857, 1–31.

Aravind, L., Abhiman, S., and Iyer, L.M. (2011). Natural history of the eukaryotic chromatin protein methylation system. Progress in molecular biology and translational science 101, 105–176.

Aravind, L., Iyer, L.M., and Koonin, E.V. (2006). Comparative genomics and structural biology of the molecular innovations of eukaryotes. Current opinion in structural biology 16, 409–419.

Aravind, L., and Koonin, E.V. (1999). DNA-binding proteins and evolution of transcription regulation in the archaea. Nucleic acids research 27, 4658–4670.

Aravind, L., and Koonin, E.V. (2000). The U box is a modified RING finger - a common domain in ubiquitination. Current biology: CB 10, R132–134.

Arnold, F.H., and Zhang, J.H. (1994). Metal-mediated protein stabilization. Trends in biotechnology 12, 189–192.

Awrey, D.E., Shimasaki, N., Koth, C., Weilbaecher, R., Olmsted, V., Kazanis, S., Shan, X., Arellano, J., Arrowsmith, C.H., Kane, C.M., et al. (1998). Yeast transcript elongation factor (TFIIS), structure and function. II: RNA polymerase binding, transcript cleavage, and read-through. The Journal of biological chemistry 273, 22595–22605.

Ball, L.J., Murzina, N.V., Broadhurst, R.W., Raine, A.R., Archer, S.J., Stott, F.J., Murzin, A.G., Singh, P.B., Domaille, P.J., and Laue, E.D. (1997). Structure of the chromatin binding (chromo) domain from mouse modifier protein 1. The EMBO journal 16, 2473–2481.

Bouazoune, K., Mitterweger, A., Langst, G., Imhof, A., Akhtar, A., Becker, P.B., and Brehm, A. (2002). The dMi-2 chromodomains are DNA binding modules important for ATP-dependent nucleosome mobilization. The EMBO journal 21, 2430–2440.

Burroughs, A.M., Ando, Y., and Aravind, L. (2014). New perspectives on the diversification of the RNA interference system: insights from comparative genomics and small RNA sequencing. Wiley interdisciplinary reviews RNA 5, 141–181.

Burroughs, A.M., Iyer, L.M., and Aravind, L. (2011). Functional diversification of the RING finger and other binuclear treble clef domains in prokaryotes and the early evolution of the ubiquitin system. Molecular bioSystems 7, 2261–2277.

Burroughs, A.M., Kaur, G., Zhang, D., and Aravind, L. (2017). Novel clades of the HU/IHF superfamily point to unexpected roles in the eukaryotic centrosome, chromosome partitioning, and biologic conflicts. Cell Cycle 16, 1093–1103.

Charier, G., Couprie, J., Alpha-Bazin, B., Meyer, V., Quemeneur, E., Guerois, R., Callebaut, I., Gilquin, B., and Zinn-Justin, S. (2004). The Tudor tandem of 53BP1: a new structural motif involved in DNA and RG-rich peptide binding. Structure 12, 1551–1562.

Chen, H.T., Legault, P., Glushka, J., Omichinski, J.G., and Scott, R.A. (2000). Structure of a (Cys3His) zinc ribbon, a ubiquitous motif in archaeal and eucaryal transcription. Protein science: a publication of the Protein Society 9, 1743–1752.

Cohen, G.B., Ren, R., and Baltimore, D. (1995). Modular binding domains in signal transduction proteins. Cell 80, 237–248.

Dalgarno, D.C., Botfield, M.C., and Rickles, R.J. (1997). SH3 domains and drug design: ligands, structure, and biological function. Biopolymers 43, 383–400.

Dillon, S.C., and Dorman, C.J. (2010). Bacterial nucleoid-associated proteins, nucleoid structure and gene expression. Nature reviews Microbiology 8, 185–195.

Doolittle, R.F. (1994). Convergent evolution: the need to be explicit. Trends Biochem Sci 19, 15–18.

Evans, P.N., Parks, D.H., Chadwick, G.L., Robbins, S.J., Orphan, V.J., Golding, S.D., and Tyson, G.W. (2015). Methane metabolism in the archaeal phylum Bathyarchaeota revealed by genome-centric metagenomics. Science 350, 434–438.

Finn, R.D., Clements, J., Arndt, W., Miller, B.L., Wheeler, T.J., Schreiber, F., Bateman, A., and Eddy, S.R. (2015). HMMER web server: 2015 update. Nucleic acids research 43, W30–38.

Finn, R.D., Clements, J., and Eddy, S.R. (2011). HMMER web server: interactive sequence similarity searching. Nucleic acids research 39, 18.

Gong, W., Wang, J., Perrett, S., and Feng, Y. (2014). Retinoblastoma-binding protein 1 has an interdigitated double Tudor domain with DNA binding activity. The Journal of biological chemistry 289, 4882–4895.

Grishin, N.V. (2000). C-terminal domains of *Escherichia coli* topoisomerase I belong to the zinc-ribbon superfamily. Journal of molecular biology 299, 1165–1177.

Grishin, N.V. (2001a). Fold change in evolution of protein structures. Journal of structural biology 134, 167–185.

Grishin, N.V. (2001b). KH domain: one motif, two folds. Nucleic acids research 29, 638–643.

Grishin, N.V. (2001c). Treble clef finger--a functionally diverse zinc-binding structural motif. Nucleic acids research 29, 1703–1714.

Guo, L., Feng, Y., Zhang, Z., Yao, H., Luo, Y., Wang, J., and Huang, L. (2008). Biochemical and structural characterization of Cren7, a novel chromatin protein conserved among Crenarchaea. Nucleic acids research 36, 1129–1137.

Holm, L., and Sander, C. (1995). Dali: a network tool for protein structure comparison. Trends Biochem Sci 20, 478–480.

Iyer, L.M., Anantharaman, V., Wolf, M.Y., and Aravind, L. (2008). Comparative genomics of transcription factors and chromatin proteins in parasitic protists and other eukaryotes. International journal for parasitology 38, 1–31.

Jones, D.O., Cowell, I.G., and Singh, P.B. (2000). Mammalian chromodomain proteins: their role in genome organisation and expression. BioEssays: news and reviews in molecular, cellular and developmental biology 22, 124–137.

Kaur, G., and Subramanian, S. (2015). The Ku–Mar zinc finger: A segment-swapped zinc ribbon in MarR-like transcription regulators related to the Ku bridge. Journal of structural biology.

Kim, D., Blus, B.J., Chandra, V., Huang, P., Rastinejad, F., and Khorasanizadeh, S. (2010). Corecognition of DNA and a methylated histone tail by the MSL3 chromodomain. Nature structural & molecular biology 17, 1027–1029.

Kishan, K.V., and Agrawal, V. (2005). SH3-like fold proteins are structurally conserved and functionally divergent. Current protein & peptide science 6, 143–150.

Klug, A. (2010). The discovery of zinc fingers and their applications in gene regulation and genome manipulation. Annual review of biochemistry 79, 213–231.

Koonin, E.V., Zhou, S., and Lucchesi, J.C. (1995). The chromo superfamily: new members, duplication of the chromo domain and possible role in delivering transcription regulators to chromatin. Nucleic acids research 23, 4229–4233.

Krishna, S.S., and Aravind, L. (2010). The bridge-region of the Ku superfamily is an atypical zinc ribbon domain. Journal of structural biology 172, 294–299.

Krishna, S.S., and Grishin, N.V. (2004). Structurally analogous proteins do exist! Structure 12, 1125–1127.

Krishna, S.S., Majumdar, I., and Grishin, N.V. (2003). Structural classification of zinc fingers: survey and summary. Nucleic acids research 31, 532–550.

Kryshtafovych, A., Moult, J., Bales, P., Bazan, J.F., Biasini, M., Burgin, A., Chen, C., Cochran, F.V., Craig, T.K., Das, R., et al. (2014). Challenging the state of the art in protein structure prediction: Highlights of experimental target structures for the 10th Critical Assessment of Techniques for Protein Structure Prediction Experiment CASP10. Proteins 82 Suppl 2, 26–42.

Kuriyan, J., and Cowburn, D. (1993). Structures of SH2 and SH3 domains: Current opinion in structural biology 1993, 3:828–837. Current opinion in structural biology 3, 828–837.

Lassmann, T., Frings, O., and Sonnhammer, E.L. (2009). Kalign2: high-performance multiple alignment of protein and nucleotide sequences allowing external features. Nucleic acids research 37, 858–865.

Lingel, A., Simon, B., Izaurralde, E., and Sattler, M. (2003). Structure and nucleic-acid binding of the Drosophila Argonaute 2 PAZ domain. Nature 426, 465–469.

Luger, K., Mader, A.W., Richmond, R.K., Sargent, D.F., and Richmond, T.J. (1997). Crystal structure of the nucleosome core particle at 2.8 A resolution. Nature 389, 251–260.

Lupas, A.N., Ponting, C.P., and Russell, R.B. (2001). On the evolution of protein folds: are similar motifs in different protein folds the result of convergence, insertion, or relics of an ancient peptide world? Journal of structural biology 134, 191–203.

Maurer-Stroh, S., Dickens, N.J., Hughes-Davies, L., Kouzarides, T., Eisenhaber, F., and Ponting, C.P. (2003). The Tudor domain 'Royal Family': Tudor, plant Agenet, Chromo, PWWP and MBT domains. Trends Biochem Sci 28, 69–74.

Moriscot, C., Gribaldo, S., Jault, J.M., Krupovic, M., Arnaud, J., Jamin, M., Schoehn, G., Forterre, P., Weissenhorn, W., and Renesto, P. (2011). Crenarchaeal CdvA forms double-helical filaments containing DNA and interacts with ESCRT-III-like CdvB. PloS one 6, e21921.

Murzin, A.G. (1998). How far divergent evolution goes in proteins. Current opinion in structural biology 8, 380–387.

Murzin, A.G. (2008). Biochemistry. Metamorphic proteins. Science 320, 1725–1726.

Murzin, A.G., Brenner, S.E., Hubbard, T., and Chothia, C. (1995). SCOP: a structural classification of proteins database for the investigation of sequences and structures. Journal of molecular biology 247, 536–540.

Nielsen, P.R., Nietlispach, D., Mott, H.R., Callaghan, J., Bannister, A., Kouzarides, T., Murzin, A.G., Murzina, N.V., and Laue, E.D. (2002). Structure of the HP1 chromodomain bound to histone H3 methylated at lysine 9. Nature 416, 103–107.

Olmsted, V.K., Awrey, D.E., Koth, C., Shan, X., Morin, P.E., Kazanis, S., Edwards, A.M., and Arrowsmith, C.H. (1998). Yeast transcript elongation factor (TFIIS), structure and function. I: NMR structural analysis of the minimal transcriptionally active region. The Journal of biological chemistry 273, 22589–22594.

Orengo, C.A., Sillitoe, I., Reeves, G., and Pearl, F.M. (2001). Review: what can structural classifications reveal about protein evolution? Journal of structural biology 134, 145–165.

Pandit, S.B., and Skolnick, J. (2008). Fr-TM-align: a new protein structural alignment method based on fragment alignments and the TM-score. BMC bioinformatics 9, 531.

Pawson, T. (1995). Protein modules and signalling networks. Nature 373, 573–580.

Ponting, C.P., Aravind, L., Schultz, J., Bork, P., and Koonin, E.V. (1999). Eukaryotic signalling domain homologues in archaea and bacteria. Ancient ancestry and horizontal gene transfer. Journal of molecular biology 289, 729–745.

Reeve, J.N., Bailey, K.A., Li, W.T., Marc, F., Sandman, K., and Soares, D.J. (2004). Archaeal histones: structures, stability and DNA binding. Biochemical Society transactions 32, 227–230.

Robinson, H., Gao, Y.G., McCrary, B.S., Edmondson, S.P., Shriver, J.W., and Wang, A.H. (1998). The hyperthermophile chromosomal protein Sac7d sharply kinks DNA. Nature 392, 202–205.

Roessler, C.G., Hall, B.M., Anderson, W.J., Ingram, W.M., Roberts, S.A., Montfort, W.R., and Cordes, M.H. (2008). Transitive homology-guided structural studies lead to discovery of Cro proteins with 40% sequence identity but different folds. Proceedings of the National Academy of Sciences of the United States of America 105, 2343–2348.

Salgado, E.N., Radford, R.J., and Tezcan, F.A. (2010). Metal-directed protein self-assembly. Accounts of chemical research 43, 661–672.

Schwede, T., and Peitsch, M.C. (2008). Computational Structural Biology: Methods and Applications (World Scientific Publishing Company).

Soding, J., Biegert, A., and Lupas, A.N. (2005). The HHpred interactive server for protein homology detection and structure prediction. Nucleic acids research 33,,W244–248.

Subramanian, G., Koonin, E.V., and Aravind, L. (2000). Comparative genome analysis of the pathogenic spirochetes Borrelia burgdorferi and Treponema pallidum. Infection and immunity 68, 1633–1648.

Swindells, M.B., Orengo, C.A., Jones, D.T., Hutchinson, E.G., and Thornton, J.M. (1998). Contemporary approaches to protein structure classification. BioEssays: news and reviews in molecular, cellular and developmental biology 20, 884–891.

Wickham, H. (2009). ggplot2: Elegant Graphics for Data Analysis (Springer-Verlag New York).

Xu, C., Cui, G., Botuyan, M.V., and Mer, G. (2015a). Methyllysine Recognition by the Royal Family Modules: Chromo, Tudor, MBT, Chromo Barrel, and PWWP Domains. In Histone Recognition, M.-M. Zhou, ed. (Cham: Springer International Publishing), pp. 49–82.

Xu, Q., Mengin-Lecreulx, D., Liu, X.W., Patin, D., Farr, C.L., Grant, J.C., Chiu, H.J., Jaroszewski, L., Knuth, M.W., Godzik, A., et al. (2015b). Insights into Substrate Specificity of NlpC/P60 Cell Wall Hydrolases Containing Bacterial SH3 Domains. mBio 6, e02327–02314.

Yan, K.S., Yan, S., Farooq, A., Han, A., Zeng, L., and Zhou, M.M. (2003). Structure and conserved RNA binding of the PAZ domain. Nature 426, 468–474.

Zhang, D., Iyer, L.M., Burroughs, A.M., and Aravind, L. (2014). Resilience of biochemical activity in protein domains in the face of structural divergence. Current opinion in structural biology 26, 92–103.

Zhang, Y., and Skolnick, J. (2005). TM-align: a protein structure alignment algorithm based on the TMscore. Nucleic acids research 33, 2302–2309.

Zhang, Z., Gong, Y., Chen, Y., Li, H., and Huang, L. (2015). Insights into the interaction between Cren7 and DNA: the role of loop beta3-beta4. Extremophiles: life under extreme conditions.

Zhang, Z., Gong, Y., Guo, L., Jiang, T., and Huang, L. (2010). Structural insights into the interaction of the crenarchaeal chromatin protein Cren7 with DNA. Molecular microbiology 76, 749–759.

Zhang, Z., Guo, L., and Huang, L. (2012). Archaeal chromatin proteins. Science China Life Sciences 55, 377–385.

